# Inhibition of the TLR4 signalling pathway with TAK-242 reduces RSV infection and cytokine release in primary airway epithelial cells

**DOI:** 10.1101/2021.11.22.469503

**Authors:** Lindsay Broadbent, Grace C. Roberts, Jonathon D. Coey, Judit Barabas, Michael D. Shields, Ultan F. Power, the Breathing Together Consortium

## Abstract

Respiratory syncytial virus (RSV) infection is the leading cause of hospitalisation in children worldwide, but there is still no vaccine or anti-viral treatment available. RSV has been implicated in the development of respiratory diseases such as asthma. Toll like receptor 4 (TLR4) has been well characterised in the immune responses to RSV. However, the role of TLR4 in RSV infection remains unclear. To study RSV in the lung epithelium, where RSV preferentially infects ciliated cells, we used a well-differentiated primary airway epithelial cell (WD-PAEC) model: a pseudostratified epithelium that produces mucus and beating cilia. We demonstrate in this physiologically relevant model that TLR4 is a pro-viral factor. Inhibition of TLR4 using TAK-242 significantly reduces RSV titres in WD-PAECs in a dose-dependent manner but has no effect on RSV growth kinetics in a range of immortalised respiratory-derived cell lines. Specific inhibition of a range of downstream effectors of TLR4 signalling in the WD-PAEC model identified p38 MAPK as a pro-viral factor, whereas inhibition of MEK1/2 significantly increased RSV titres. Our data demonstrate a role for TLR4 in RSV infection and highlight the importance of biologically relevant models to study virus-host interactions.

**Author summary:** Respiratory Syncytial Virus (RSV) can cause severe respiratory infection in young children and is responsible for approximately 200,000 deaths worldwide every year. Despite decades of research since the identification of this virus in the 1950s there is still no vaccine or treatment available. Advances in research have led to the development of cell cultures that are very similar to the cells that line human airways. These cultures provide an opportunity to study how viruses interacts with airway cells in a representative model and may provide insights that traditional research models have not yet been able to answer. Using this experimental model we show that a drug, TAK-242 which targets a pathogen recognition receptor on the surface of cells, reduces growth of RSV and dampens the immune response to infection in these airway cells. Our data demonstrate potential targets for RSV treatments and also highlight the importance of using relevant experimental models.

## Introduction

Respiratory syncytial virus (RSV) is responsible for over 3 million hospitalisations of infants each year and is the primary cause of severe lower respiratory tract infections (LRTIs) in children worldwide (1,2). There are currently no available vaccines or effective therapeutics against RSV.

As with many viruses, the innate immune system plays a critical role in defence against RSV disease. Innate immune signalling pathways are triggered by the detection of pathogen associated molecular patterns (PAMPs) by surface and/or intracellular pathogen recognition receptors (PRRs). For RNA viruses, toll-like receptor 3 (TLR3) has been well characterised as a PRR that detects dsRNA and induces activation of IFN regulatory factor 3 (IRF3) and the NF-κB pathway (3). TLR4 is also able to activate IRF3 and NF-κB pathways but is better characterised as a receptor of lipopolysaccharide (LPS), a PAMP produced by many Gram-negative bacterial species. TLR4 is part of the LPS receptor complex, alongside myeloid differentiation 2 (MD2), both of which are expressed on the surface of cells from myeloid and epithelial cells (4). In addition to LPS, several viral proteins have been identified as ligands for TLR4, including Ebola virus glycoprotein (5) and RSV fusion (F) protein (6–8). The RSV F protein interacts with the TLR4 complex, initiating a signalling cascade leading to the production of pro-inflammatory cytokines (9). Furthermore, a functional TLR4 is required for NF-κB activation in both cell lines and mice (10).

TLR4 expression increases in cells infected with RSV (11,12). An increased expression of a PRR would be expected upon viral infection. Several studies have also indicated that TLR4 enhances RSV production. Previous studies using HEK293T cells have highlighted a role for TLR4 in the RSV lifecycle, through overexpression and knockdown of TLR4 (13,14).

An issue perhaps causing confusion in the literature is an appropriate choice of model – both cellular and viral. Many studies have used mouse models, which have a different TLR4 to humans, both in terms of the protein structure and its ligands (15). Many of the studies discussed above used immortalised cell lines in 2D culture which, although incredibly versatile, are limited in representing the complexity of the airway epithelium (16). In addition, research into TLR4-RSV interactions commonly use the prototypic A2 strain of RSV, which is considered to be cell culture-adapted with several mutations, including within the F protein (17).

In the clinic, TLR4 has been linked to severe RSV disease (18). TLR4 SNPs, Asp299Gly and Thr399Ile were found to be over-represented in infants hospitalised with RSV bronchiolitis (19). Substantial increases of TLR4 expression in peripheral blood monocytes were also linked to severe RSV disease and to low oxygen saturation (20). In addition, reduced levels of miRNA-140-5p, which targets TLR4, were found in the nasopharynx and peripheral blood of patients with severe RSV disease (21).

In our laboratory, we are particularly interested in the interaction between RSV and the airway epithelium. Therefore, we use well-differentiated primary airway epithelial cells (WD-PAECs) cultured in an air-liquid interface (ALI) resulting in a pseudostratified epithelium that closely resembles that of airway tissue *in vivo* to study interactions between the airway epithelium and RSV.

Here, we demonstrated that inhibition of TLR4 greatly reduces RSV viral growth kinetics in the physiologically relevant WD-PAEC model and that this effect was not observed in immortalised airway epithelial cell lines. We demonstrated that downstream effectors of TLR4 signalling, including p38 MAPK and NF-κB, were involved in RSV replication kinetics and IFNλ1 secretion, respectively. These results highlight the importance of using appropriate models to investigate, or validate, host-pathogen interactions.

## Results

### Efficient RSV infection is dependent on TLR4 expression in HEK293T cells

To determine whether TLR4 plays a role in RSV infection, HEK293T cells, either overexpressing TLR4 (through stable transfection) or TLR4-null cells, were infected with RSV-A2-GFP. GFP fluorescence was assessed to determine the kinetics of spread of RSV infection. Significantly higher levels of infection were observed in the HEK293/TLR4 cells compared to the HEK293/null cells, indicating that TLR4 is important in the RSV lifecycle (Figure 1).

**Figure 1.**
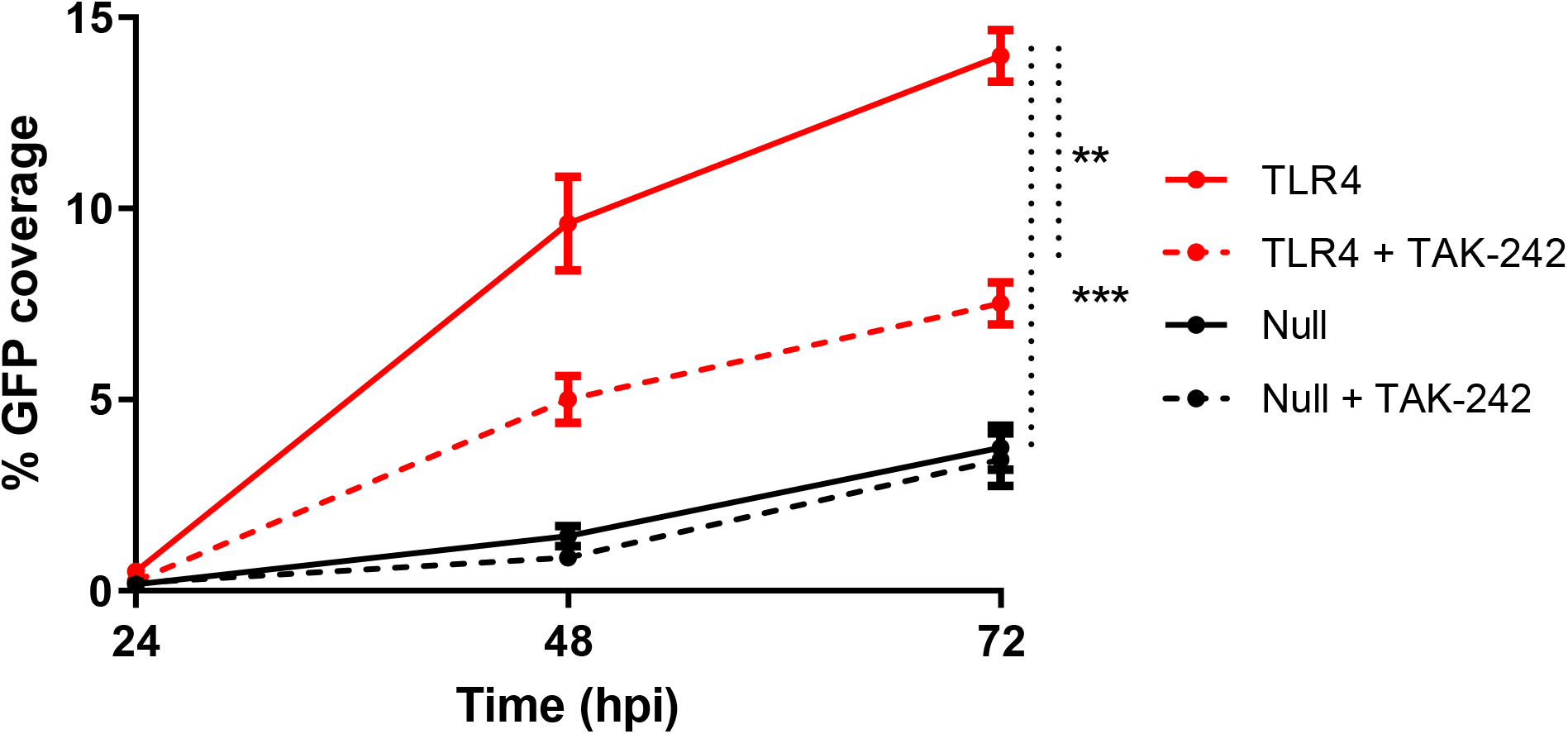
RSV replicates well in TLR4-expressing cells but poorly in TLR4-null cells. HEK293/null and HEK293/TLR4 cells were treated with 10 μg/mL TAK-242 or DMSO (mock) for 6 h, infected with RSV-A2-GFP at MOI=0.1. Five images per well were captured at 24, 48, and 72 hpi using a Nikon TE2000U microscope. The percentage of the image expressing green fluorescence was assessed using Image J. (Data shown is n=4 from two independent experiments. Areas under the curves for each condition were calculated and significance determined by paired T-test where ** = p<0.01, *** = p<0.001).

To confirm the role of TLR4 in the RSV lifecycle, a TLR4 inhibitor, TAK-242 (also known as CLI-095 or Resatorvid) was used. TAK-242 is a small molecule inhibitor that binds to the intracellular domain of TLR4 and inhibits downstream signalling, but does not affect ligand binding (22). Both the HEK293/TLR4 and HEK293/null cells were treated with TAK-242 prior to RSV infection. In the TLR4-overexpressing cells, treatment significantly reduced RSV replication, while it had no effect in the TLR4 null cells, indicating that TLR4 signalling is important for RSV infection.

### Inhibition of TLR4 or MD2 has no effect on RSV growth kinetics in immortalised respiratory-derived cell lines

Prior to conducting experiments on cell lines, the expression of TLR4 and MD2 was confirmed through immunofluorescence (IF). A range of human respiratory epithelium-derived cell lines, including A549 (adenocarcinoma alveolar epithelial cells), BEAS-2B (bronchial epithelial cells), and Calu-3 (lung epithelial adenocarcinoma) cells were either mock infected or infected with the low passage clinical isolate RSV-BT2a (MOI=0.1) for 24 h and stained for TLR4 and MD2 (Figure 2A). For all cell lines, TLR4 was predominantly located in the cell membrane, whereas MD2 appeared more diffuse throughout the cytoplasm. When infected with RSV-BT2a, both TLR4 and MD2 expression appeared to be increased in A549 and BEAS-2B cells, but not in Calu-3 cells.

**Figure 2.**
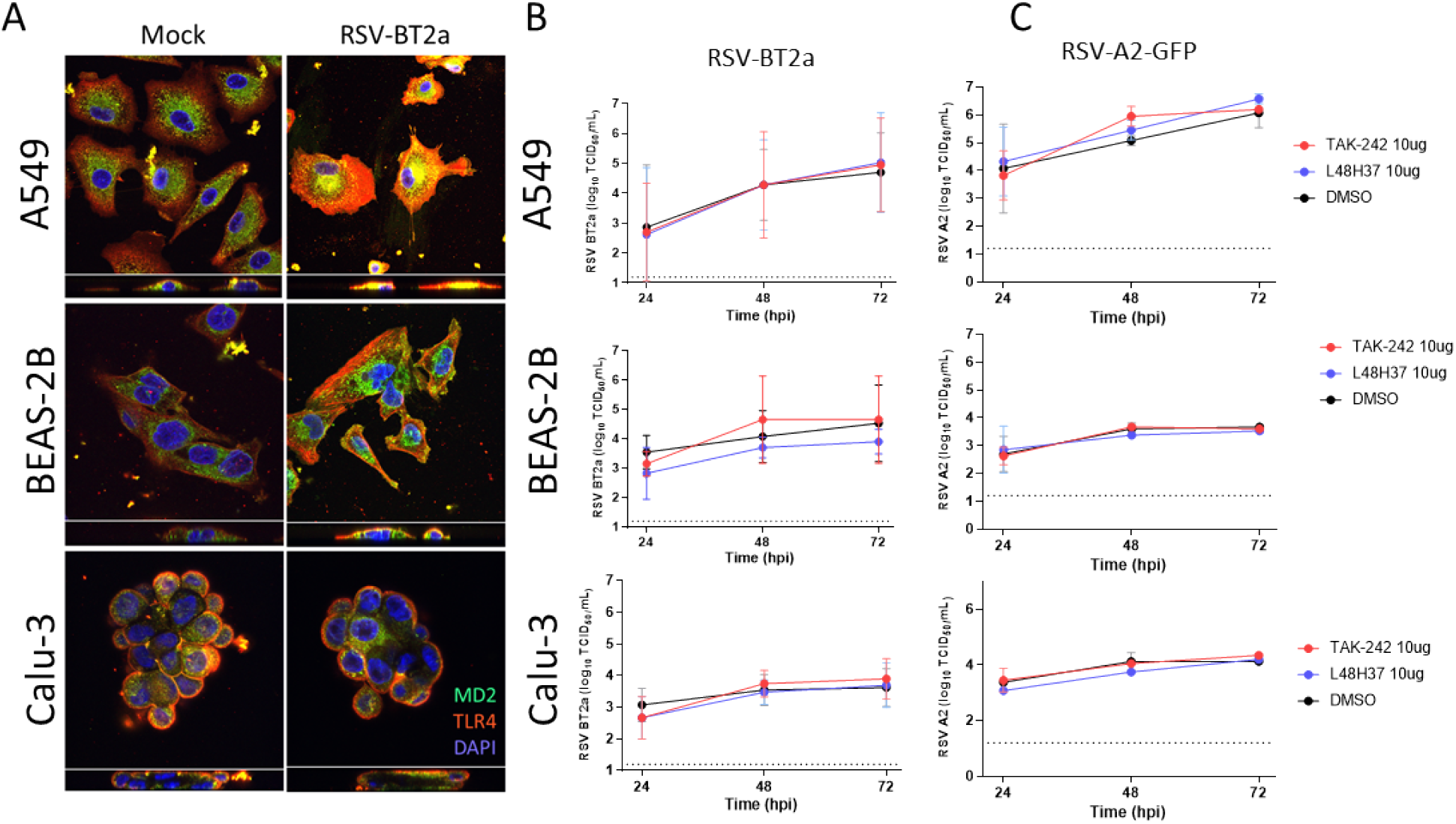
Inhibition of TLR4 or MD2 had no effect on RSV production in respiratory-derived immortalised cell lines. (A) A549, BEAS-2B, and Calu-3 cells were either mock or RSV-BT2a infected (MOI=0.1), fixed at 24 hpi, and stained for TLR4 (red) and MD2 (green) to confirm expression of both proteins (images taken via a Leica SP5 confocal microscope at 63x magnification). (B) A549, BEAS-2B, and Calu-3 cells were pre-treated with 10 ug/mL of either TAK-242 or L48H37, or DMSO (mock) for 6h prior to infection with RSV-BT2a at MOI=0.1. Media was harvested at 24, 48 and 72 hpi and RSV titre quantified via TCID_50_ assay. (C) As described in part B, but infected with RSV-A2-GFP.

These cell lines were then treated with either TAK-242 (TLR4 inhibitor) or L48H37 (an MD2 inhibitor) at 10 μg/mL 6 h prior to infection with RSV BT2a or RSV A2-GFP. There was no effect on virus titres throughout the 72 h time course for either RSV BT2a (Figure 2B) or RSV A2-GFP (Figure 2C). These data are consistent with previous reports that found little evidence that TLR4 played a significant role in RSV pathogenesis in immortalised cell lines (14).

### Removal of heparan sulphate from A549 cells reduces RSV-A2 but not RSV-BT2a viral titres and has no impact on the effect of TAK-242 treatment

Due to the abundance of heparan sulfate (HS) on immortalised cell lines, we hypothesised that RSV may be primarily utilising HS for cell entry, masking any potential effect of TLR4 inhibition. HS was shown to facilitate entry of RSV into immortalised cells (23–25). However, there is debate in the literature as to whether HS is truly involved in RSV entry *in vivo,* as ciliated human airway epithelial cells, which RSV preferentially infect, express no or limited amounts of HS (26).

To assess whether HS was masking any effect of TLR4 inhibition in cell lines, A549 cells were treated with TAK-242 and/or treated with heparinase (to remove cell surface HS) prior to infection with RSV (either BT2a or A2-GFP). There was no significant reduction in RSV-BT2a titres following TAK-242, heparinase, or the combination treatment (Figure 3A). In contrast, RSV-A2-GFP growth kinetics, as measured by GFP expression, were significantly reduced following heparinase treatment and the combination TAK-242 and heparinase treatment, but not following TAK-242 alone (Figures 3B and 3C). Viral titrations were consistent with a reduction in RSV-A2-GFP titres in these conditions (Figure 3A), but the results did not reach significance.

**Figure 3.**
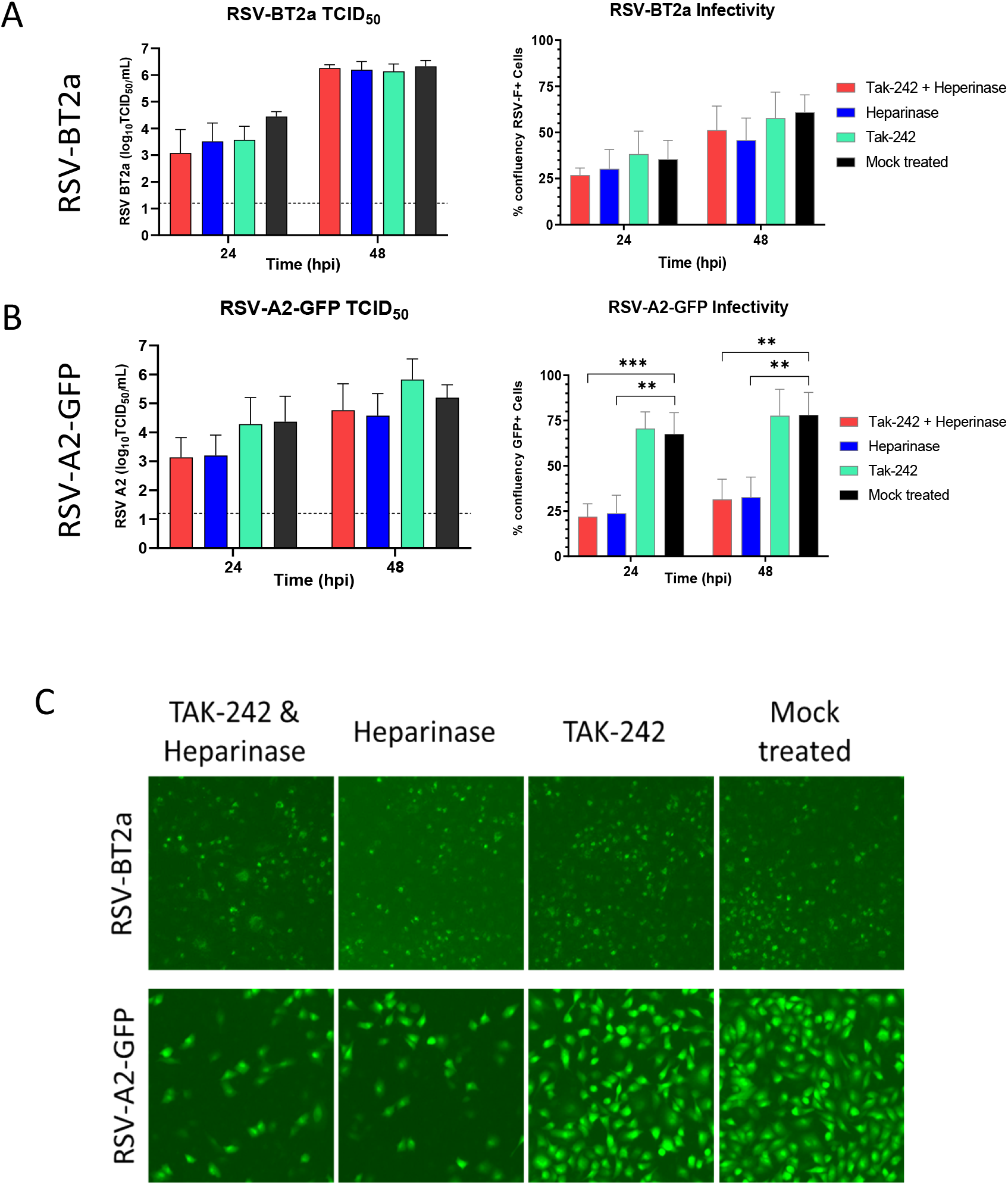
Pre-treating A549 cells with heparinase in combination with TAK-242 significantly reduced the number of infected cells, but not RSV production. A549 cells were treated with either 10 μg/mL TAK-242 or DMSO (mock) for 5 h, then heparinase-I-(5 mIU/mL) or PBS-(mock) treated for 1 h. Cells were then infected with RSV-BT2a (A) or RSV-A2-GFP (B) at an MOI=1 for 2 h. Supernatant was collected from cells at 24 and 48 hpi and titred via TCID_50_ assay (n=4). Corresponding cells were then fixed (BT2a-infected cells stained for RSV-F) and scanned for brightfield and green fluorescence using a Celigo Imaging Cytometer (Nexcelom). Celigo software was used to quantify number of infected cells by quantifying % confluence of fluorescence cells, and % confluence total cells. Data shown is green confluency normalised to total cell confluency (n=4). (C) Representative images of Celligo-scanned wells at 24 hpi.

### Inhibition of TLR4 significantly reduces RSV replication and pro-inflammatory responses in WD-PAEC cultures

The use of primary airway epithelial cells, isolated from human respiratory tracts, to form well-differentiated 3D cultures is well established for the study of RSV (27–29). Culture in an air-liquid interface (ALI) promotes the development of a pseudostratified epithelium with basal cells, ciliated cells, and mucus-producing goblet cells, representative of the respiratory airway epithelium *in vivo.* We exploited these physiologically and morphologically relevant models to investigate whether TLR4 and/or MD2 played a role in RSV replication in human airway epithelium.

Expression of both TLR4 and MD2 was confirmed in WD-PAECs by IF. Fully differentiated WD-PAECs that exhibited good cilia and mucus coverage were selected and infected with RSV (MOI=0.1) or mock infected for 96 h. Cells were then fixed and stained for RSV F, TLR4, and MD2. Expression of both TLR4 and MD2 in WD-PAECs was predominately apical, as shown by the orthogonal images (Figure 4a). Following infection TLR4 expression was increased, and both TLR4 and MD2 appeared concentrated in areas proximal to RSV infection (as indicated by RSV-F staining).

**Figure 4.**
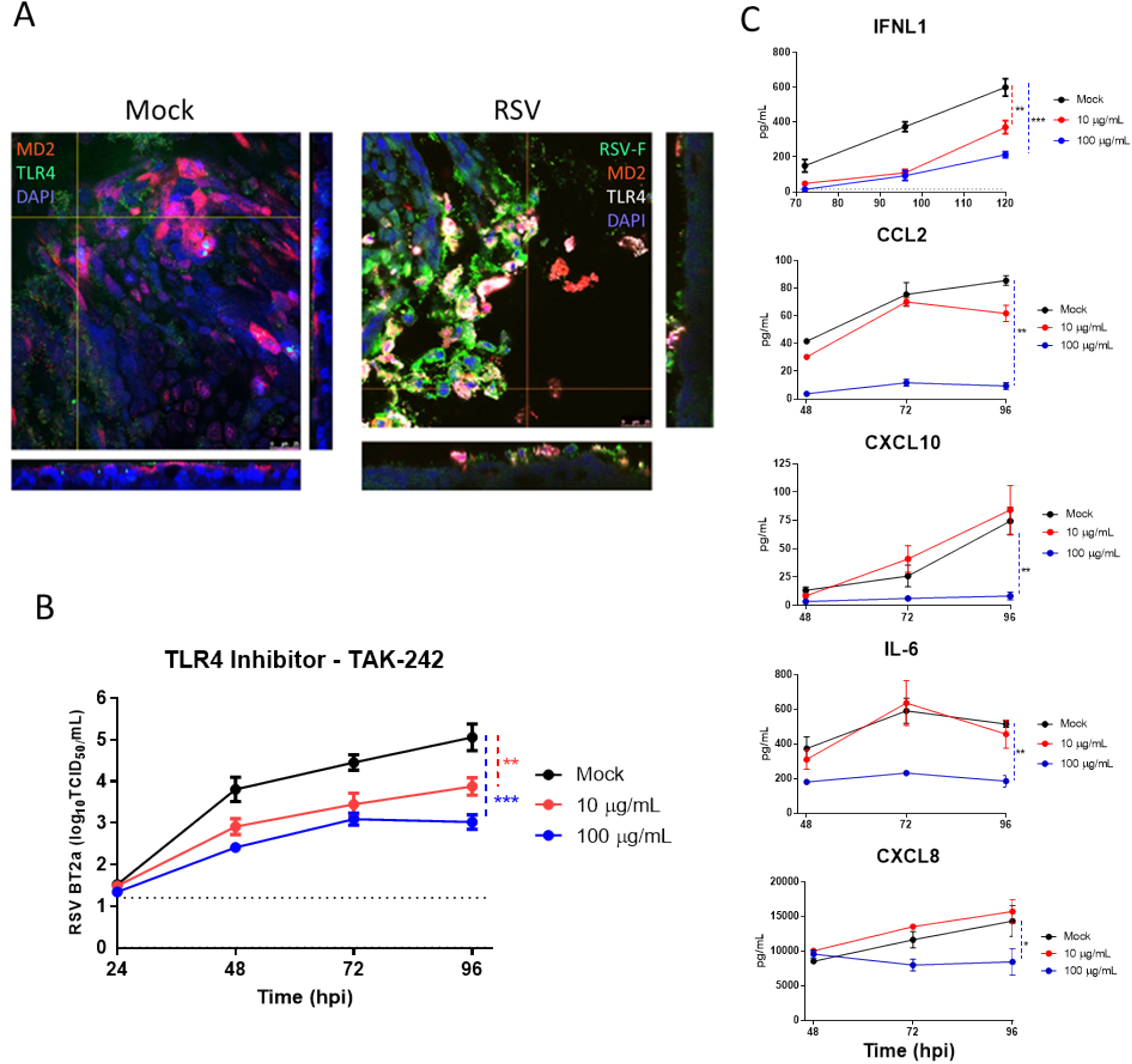
Inhibition of TLR4 significantly reduces RSV production in WD-PAECs. (A) WD-PAEC were infected with RSV-BT2a or mock infected. Cultures were fixed at 96 hpi and stained for MD2 (red), TLR4 (green) and DAPI in the left (mock) panel, and for RSV-F (green), MD2 (red), TLR4 (white) or DAPI in the right (infected) panel. (B) WD-PAECs (n=7) were apically treated with either DMSO (mock), 10 or 100 ug/mL of TAK-242 6 h prior to infection with RSV-BT2a. Apical washes were taken every 24 hpi and RSV titres quantified via TCID_50_ assay (statistical significance was calculated using a paired T test for each time point comparing treated cells to mock, where * = p<0.5, ** = p<0.01, *** = p<0.001). (C) WD-PAECs (n=3 donors) treated and infected as described in part A. Basolateral medium was harvested and replaced every 24 hpi. The concentration of IFNL1 was quantified via ELISA. CCl2, CXCL10, IL-6 and CXCL8 were quantified by BioPlex analysis (Bio-Rad). Vertical dotted lines indicate statistical significance when areas under the curves were calculated and compared to mock via T test (* = p<0.05; ** = p<0.01)

To investigate the role of TLR4 in RSV infection of WD-PAECs, cultures were pre-treated with 10 or 100 μg/mL TAK-242 for 6 h prior to infection with RSV. Apical washes were taken every 24 h over the course of a 96 h period, and RSV quantified by TCID_50_ assay. Both TAK-242 doses significantly reduced RSV titres, in a dose-dependent manner from 48 to 96 hpi, compared to the mock treated cells (Figure 4B). Furthermore, TAK-242 (100 μg/mL) significantly reduced secretion of IFNλ1 (IL-29), CCL2 (MCP-1), CXCL10 (IP-10), IL-6 and CXCL8 (IL-8) from RSV-infected WD-PAECS (Figure 4C).

### Neither inhibition of MD2 nor removal of cell-surface heparan sulfate significantly impact RSV replication in WD-PAEC cultures

WD-PAECs were treated with the MD2 inhibitor L48H37 for 6 h prior to infection with RSV. There was a modest reduction in RSV titres from 24-96 hpi, but this was not statistically significant (Figure 5A). L48H37 binds to the binding pocket of MD2, which was shown to prevent LPS binding and subsequent signal transduction (30). Structural modelling suggests that L48H37 does not affect MD2 from interacting with TLR4.

**Figure 5.**
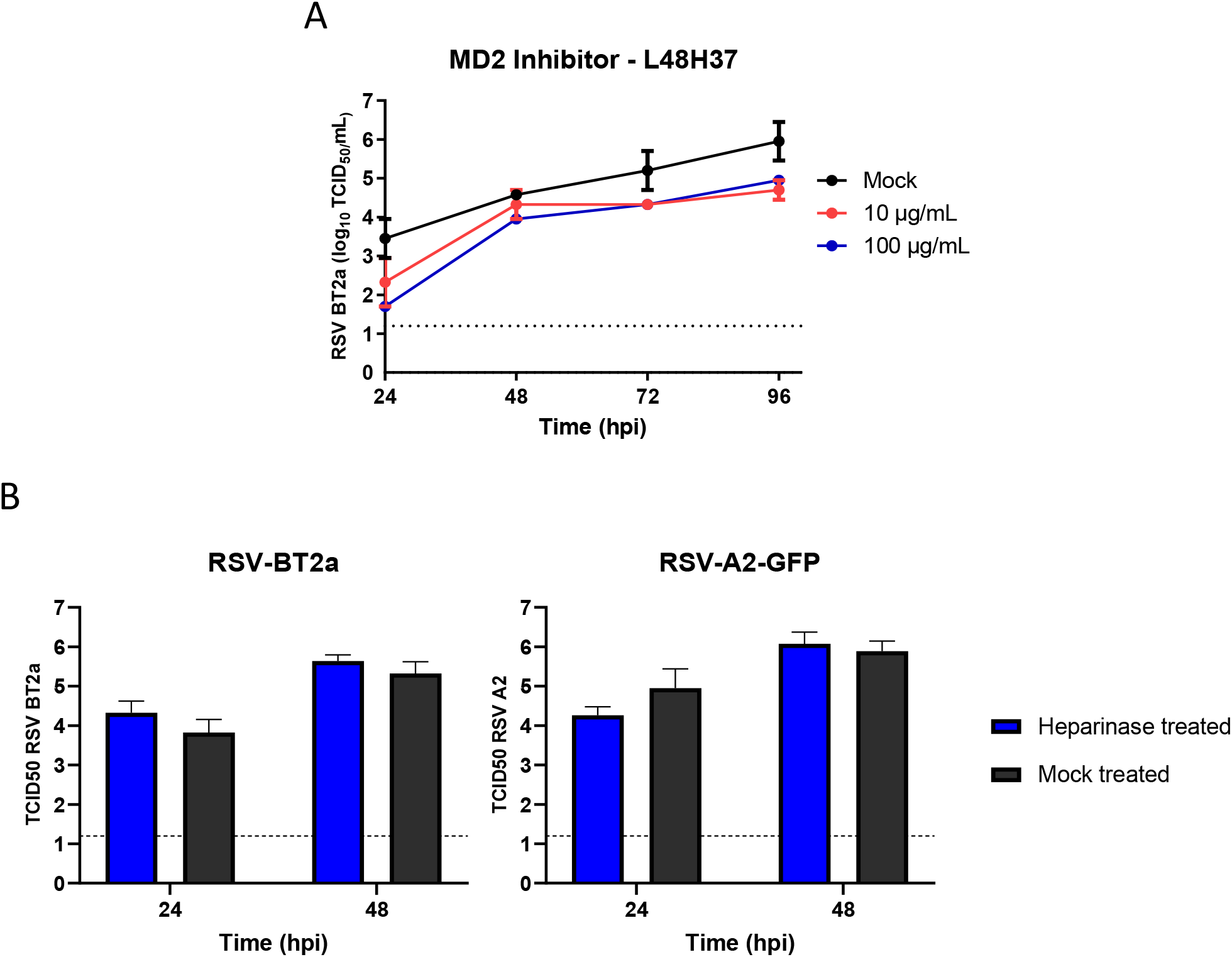
Neither inhibition of MD2, nor removal of heparin sulfate significantly affect RSV replication in WD-PAECs. (A) WD-PAECs (n=2) were apically treated with either DMSO (mock), 10 or 100 ug/mL L48H37 (MD2 inhibitor) for 6 h prior to infection with RSV-BT2a. Apical washes were taken every 24 hpi and RSV titres quantified via TCID_50_ assay. (B) WD-PAECs (n=4) were treated with heparinase-I (5 mIU/mL) or PBS (mock) for 1 h. Cells were then infected with RSV-BT2a or RSV-A2-GFP for 2 h. Apical washes were collected from cells at 24 and 48 hpi and quantified via TCID_50_ assay.

Similarly, when WD-PAECs were treated for 1 h with heparinase prior to RSV infection, there was no significant difference in viral production for either RSV-BT2a or RSV-A2-GFP (Figure 5B).

### Inhibition of TLR4 pathway intermediates identified pro- and anti-viral effectors

When activated, TLR4 triggers several downstream pathways, as detailed in Figure 6A. To determine whether the TAK-242-mediated reduction of RSV production was due to the inhibition of TLR4 itself, or the inhibition of one of the downstream effectors, WD-PAECs were treated with a range of specific inhibitors for PI3K (LY294002), MEK1/2 (U0126), p38 MAPK (SB203580), and NF-κB (JSH-230) prior to infection with RSV. Apical washes were harvested every 24 h for a 96 h period and infectious virus was quantified by TCID_50_ assay. In parallel, basal secretion of IFNλ1 was quantified via ELISA (Figure 6C). Type III IFNs, specifically IFNλ1, have been shown to be the major IFN produced in response to RSV infection in WD-PAEC models (31,32).

**Figure 6.**
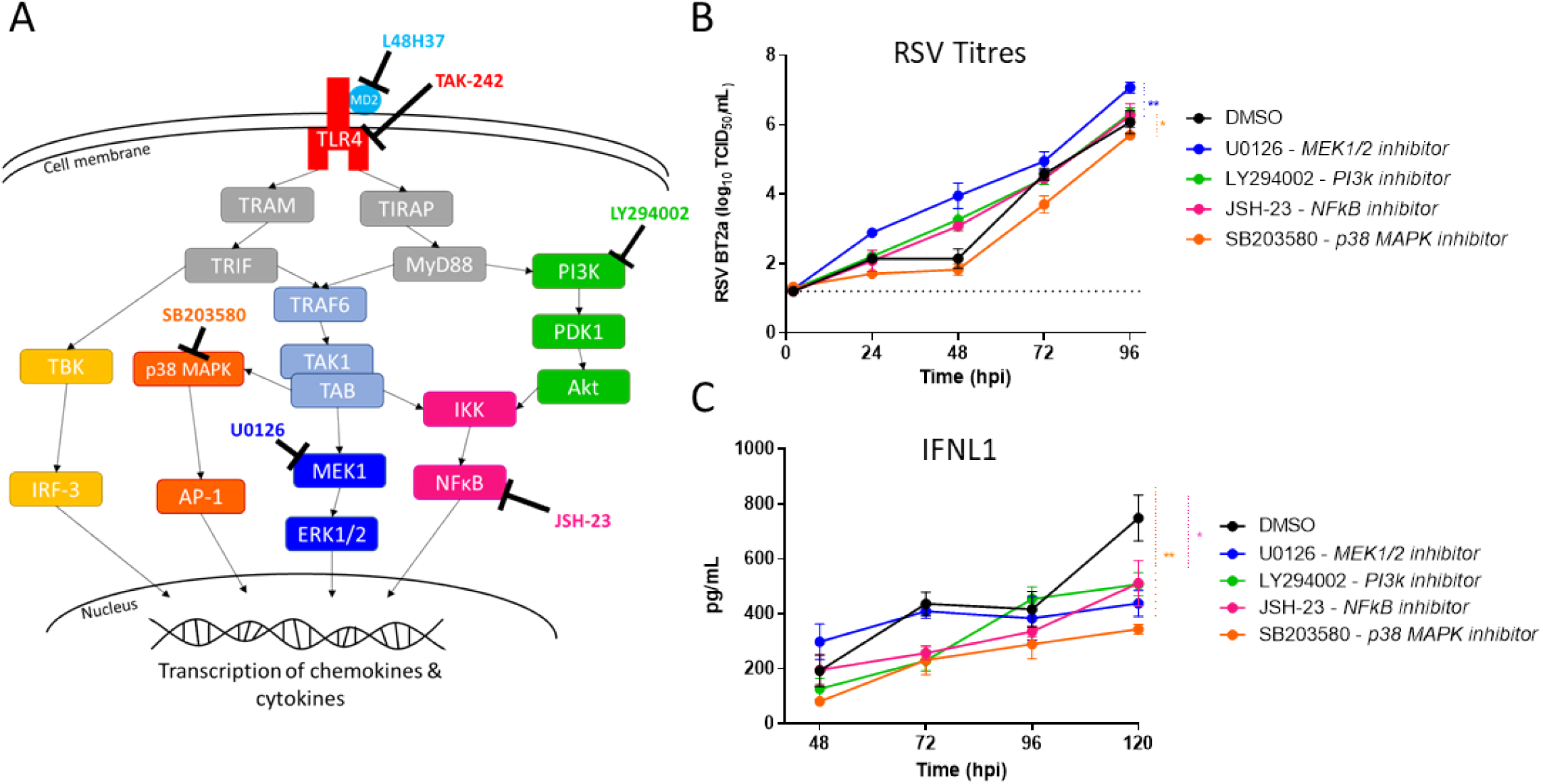
Inhibition of TLR4 downstream pathways elicited different effects on RSV production, and most reduced IFNλ1 secretion of RSV-infected WD-PAECs. (A) Diagram of downstream TLR4 signalling with the specific inhibitors used in part B indicated. (B) WD-PAECs (n=3 donors) were apically pre-treated with 50 μM LY294002 (PI3K inhibitor), 20 μM U0126 (MEK1/2 inhibitor), 1 μM SB203580 (p38 MAPK inhibitor), 20 μM JSH-23 (NF-κB inhibitor), or DMSO (mock) for 1 h at 37°C prior to infection with RSV-BT2a at MOI=0.1. Apical washes were harvested every 24 h following infection and RSV titres quantified via TCID_50_ assay. (C) Corresponding basolateral medium was harvested and replaced every 24 hpi. IFNL1 in basolateral medium was quantified by ELISA. Area under the curve was calculated for each condition and statistical significance determined via paired T test comparing each condition to the mock (* = p <0.5, **= p<0.01).

Inhibition of p38 MAPK (SB203580) significantly reduced RSV titres over the 96 h period compared to mock-treated controls, although this was not as marked a reduction as observed following TLR4 inhibition via TAK-242 (Figure 4B). IFNλ1 secretion was also reduced following p38 MAPK inhibition, possibly due to the reduction in viral growth kinetics. Conversely, inhibition of MEK1/2 significantly increased RSV titres but did not result in concomitant increases in IFNλ1 secretion. Interestingly, inhibition of NF-κB with JSH-23 significantly reduced IFNλ1 secretion but did not affect viral titres, indicating that NF-κB is implicated in RSV induction of type III IFNs.

## Discussion

Here we demonstrate that TLR4 inhibition via TAK-242 reduces RSV titres in the WD-PAEC cell model, but not in immortalised cell lines. TLR4 was previously implicated in RSV biology with studies demonstrating the role of TLR4 in effective immune responses to RSV (7,9,10,14). However, there have been conflicting reports on the involvement of TLR4 in RSV infection (11,13,14), likely caused by differences in experimental designs. We employed the use of WD-PAECs, together with a low passage clinical isolate of RSV, to assess the role of TLR4 in a physiologically relevant model of RSV infection of human airway epithelial cells.

TAK-242 demonstrated substantial prophylactic properties against RSV infection in our WD-PAEC model. As the inhibitor acts by blocking intracellular signalling and does not interfere with the extra-cellular domain of TLR4, it is unlikely that TAK-242 interferes with the RSV F/TLR4 interaction. Indeed, Ii et al demonstrated that TAK-242 does not interfere with the binding of LPS to TLR4 (33). As such, diminished RSV growth kinetics are unlikely to be due to the blocking of RSV F protein binding to the TLR4 complex. An association between TLR4 and nucleolin, a putative RSV co-receptor or entry factor, was recently described (8). Further work is needed to establish whether TAK-242 impacted the ability of RSV to interact with nucleolin and, thereby, the ability of RSV to infect epithelial cells.

Interestingly, TLR4 antagonists have been associated with a reduction in titres of other viruses, both *in vitro* and *in vivo*. Eritoran, which binds to MD2 and prevents TLR4 activation, decreased viral titres, cytokine production, clinical symptoms, and morbidity in a mouse model of influenza virus infection (34). When using MD2 inhibitor L48H37 in the WD-PAEC model, there was a slight reduction in RSV titres but this was not statistically significant. Both L48H37 and Eritoran are known to directly bind the hydrophobic pocket of MD2 and prevent ligands binding (30,35).

As TLR4 inhibition significantly reduced RSV titres release from WD-PAECs, whereas MD2 inhibition did not, this implies that TLR4 alone is important for efficient RSV infection, rather than the receptor complex as a whole. As TAK-242 specifically inhibits TLR4 signalling and not extracellular ligand binding, it is likely that a downstream effector function of TLR4 is required for RSV infection and/or replication.

Unlike our RSV/WD-PAEC model, cell lines do not replicate the morphological or physiological complexities of differentiated airway epithelium, nor the cytopathogenesis of RSV in airway epithelium *in vivo*. Indeed, even the mechanism of viral entry may differ between the two experimental models. Previous work demonstrated the role of heparan sulfate in RSV infection of continuous cell lines (23,36,37). Enzymatic removal of heparan sulfate from A549 cells greatly reduced RSV infection when using the lab-adapted RSV-A2 strain, but did not impact the results of TAK-242 treatment, indicating that even when HS is removed from the cell surface the mechanism of viral infection differs between cell lines and primary airway epithelial cells. Our data is consistent with previous studies that show the lab-adapted RSV strain A2 is heavily reliant on HS for cell entry and, therefore, may not be the most appropriate model virus for the study of the RSV lifecycle.

Used throughout this study, RSV-BT2a is a low passage clinical isolate, which, alongside many other clinical isolates, demonstrated differences in cytopathogenicity, viral growth kinetics, and pro-inflammatory responses in both primary monolayer airway epithelium and ALI cultures compared to the laboratory adapted RSV A2 strain (29). Unlike RSV A2, in this study the removal of heparan sulfate had no effect on resulting RSV BT2a titres or infectivity, suggesting that this strain is likely a more relevant virus model to study RSV replication and pathogenesis.

In humans, TLR4 SNPs and increased levels of TLR4-specific siRNA have been linked to severe RSV disease (19,21). TLR4 has been implicated in other inflammatory diseases and TAK-242 has been investigated as a therapeutic for many of these conditions, such as sepsis, brain injury, and several different types of cancer (38–42).

Following inhibition of a range of proteins downstream of TLR4 signalling, only p38-MAPK inhibition significantly reduced RSV titres. This is consistent with other reports, which have shown that p38 MAPK is activated within 10 min of RSV infection via the TLR4 adaptor protein MyD88 (13). It is intriguing, however, that when we inhibited p38 MAPK using SB203580, RSV titres were reduced to a lesser extent compared to the reduction resulting from TLR4 inhibition. This indicates that either TLR4 itself is involved in the RSV lifecycle and/or TLR4 induces multiple downstream effectors, in addition to p38 MAPK, that are required for efficient RSV infection/replication that are yet to be characterised. Intriguingly, inhibition of MEK1 and 2 significantly increased RSV titres, indicating that some aspects of the TLR4 signalling cascade are antiviral. Other studies have found inhibition of MEK1/2 to restrict RSV production in A549 cells. However, this was only evident when MEK1/2 were inhibited at 4 h post infection, indicating that MEK1/2 has a temporally-specific role in the RSV lifecycle (43). Another study, using undifferentiated AECs and RSV-A2 found that pre-treatment of cells with a MEK1/2 inhibitor did not affect RSV genome copy numbers at 24 hpi (44). The role of MEK1/2 in RSV infection requires further investigation.

In terms of a mechanism, both our data, and previous reports indicate a role for TLR4 at an early stage of the RSV lifecycle. It has been shown that both RSV F binding of TLR4, and p38 MAPK activation occur within 10 min of addition of virus to cells (6,13). Previous studies have produced conflicting data regarding the role of TLR4 in RSV infection in continuous cell lines. Our use of a physiologically relevant primary airway epithelial cell model, and the use of a clinical isolate of RSV strengthens the confidence in our findings that TLR4 is important in the RSV lifecycle. However, further study is required to determine the precise role of TLR4 in the RSV lifecycle.

In summary, we have demonstrated that TLR4 is important for efficient RSV infection/replication in airway epithelium using the WD-PAEC model. Inhibition of TLR4 dramatically reduces RSV titres and appears to be acting at an early stage of RSV infection. A precise mechanism for TLR4 in the RSV lifecycle is yet to be determined. However, in light of our findings, TLR4 presents an attractive opportunity as a pharmacological target for treatment of RSV.

## Methods

### Primary cells and WD-PAEC culture

Primary nasal or bronchial epithelial cells were obtained from commercial sources (Lonza) or from fully consenting, healthy volunteers (IRAS ID 199053; London – City and East Research Ethics Committee) and developed into well-differentiated primary airway epithelial cell (WD-PAEC) cultures as previously described in detail by (27). Previously, WD-PAECs derived from nasal cells have been demonstrated to be as representative of the airway epithelium as those derived from bronchial cells (45). In brief, cells retrieved from previously frozen stocks were grown to confluency then plated into 6.5 mm, 0.4 μm pore Transwells (Corning) at a density of 3 × 10^4^ cells per Transwell. Once cells reached confluency in, air-liquid interface (ALI) culture was established by removing apical medium. After approximately 21 days of ALI culture, differentiated cells were selected for experiments after demonstrating good cilia coverage and mucus production with no holes present in the epithelium.

### Cell lines and culture

HEK293/TLR4 and HEK293/null cells (both Invitrogen) were maintained in 4.5 g/L glucose DMEM (Gibco) supplemented with 10 units/mL pen-strep (Gibco), and 5% foetal bovine serum (FBS, Gibco). A549 cells and HEp-2 cells were maintained in 4.5 g/L glucose DMEM with 10 units/mL pen-strep, and 5% FBS. BEAS-2B cells were maintained in 1 g/L (low) glucose DMEM with 10 units/mL pen-strep, and 5% FBS. Calu-3 cells were maintained in MEM (Gibco) 10 units/mL pen-strep, 10% FBS, and 1% L-glutamine.

### Viruses and titration

Isolation and characterisation of the clinical isolate RSV-BT2a was previously described (29). RSV A2-GFP was a kindly provided by Prof. Ralph Tripp (University of Georgia) and Prof. Michael Teng (University of South Florida). Infections were performed at MOIs indicated in specific figure legends by applying virus suspension either directly to cells (in the case of cell lines) or to the apical compartment of WD-PAEC cultures.

Virus titration was performed via TCID_50_ assay on HEp-2 (kindly provided by Prof. Ralph Tripp, University of Georgia) cells, as previously described (46).

### Immunofluorescence/fluorescent imaging

For imaging of cell lines, cells were cultured on coverslips and fixed for 1 h in 4% PFA (w/v in PBS). WD-PAECs were fixed by submersion in 4% PFA for 2 h. Cells were permeabilised with 0.1% Triton-X-100 (v/v in PBS, 1 h for cell lines or 2 h for WD-PAEC cultures) and blocked for 1 h in 1% BSA (w/v in PBS). Immunofluorescent staining was performed for TLR4 (1:100 dilution, Ab22048, Abcam), MD2 (1:100 dilution, Ab24182, Abcam), and RSV-F (1:500 dilution, conjugated to a 488 fluorophore MAB8262X, Merck), incubating overnight at 4°C. Appropriate secondary antibodies (AlexaFluor, Invitrogen) were applied for 1 h at 37°C prior to mounting onto slides using mounting media containing DAPI (Vectashield, Vector Labs). Images were obtained on a Leica SP5 confocal microscope or a Nexcelom Celigo as described in individual figure legends.

### Inhibitors

TAK-242 (aka CLI-095, Resatorvid) and L48H37 (both Merck) were reconstituted in DMSO at a concentration of 1 mg/mL and were further diluted in DMEM (with no supplements) for working concentrations of either 10 or 100 μg/mL. Inhibitors were applied to cells for 6 h at 37 °C.

Other signalling inhibitors were reconstituted in DMSO as per manufacturer’s instructions, and further diluted in DMEM (with no supplements) to working concentrations of double the IC50, as indicated by the manufacturer; LY294002 (50 μM, LC Labs), U0129 (20 μM, LC Labs), SB203580 (1 μM, Sigma Aldrich) and JSH-23 (20 μM, Sigma Aldrich).

### Heparinase treatment

Heparinase I (Sigma Aldrich) was reconstituted as per manufacturer’s instructions. Heparinase was resuspended in DMEM (with no supplements) at a working concentration of 5 mIU/mL prior to application to cells, which were incubated for 1 h at 37 °C.

### Quantification of cytokines and chemokines

Concentrations of IFNλ1 from basolateral medium of WD-PAECs was determined using the Platinum Sandwich ELISA kit (Invitrogen) following the manufacturer’s protocol. Concentrations of CCL2, CXCL10, IL-6 and CXCL8 were determined using the ProcartaPlex kit (Invitrogen) following the manufacturer’s protocol.

### Statistical analysis

GraphPad Prism^®^ was used to create graphical representations of the data and for statistical analyses. Summary measures over time were compared by calculating the areas under the curves (AUC). Statistical significance was determined using t tests comparing AUCs. (* p value = <0.05; ** p value = <0.01; *** p value = <0.001.)

## Acknowledgements

The authors would like to thank the donors of the primary airway cells. We would also like to thank the technical staff at the WWIEM: Nuala McCann, Mervyn McCaigue, Barney O'Loughlin, William Houppy, and Cathy Fenning.

